# Characterization of auditory responsive neurons in the mouse superior colliculus to naturalistic sounds

**DOI:** 10.1101/2025.01.13.632849

**Authors:** Yufei Si, Shinya Ito, Alan M. Litke, David A. Feldheim

## Abstract

Locating the source of a specific sound in a complex environment and determining its saliency is critical for survival. The superior colliculus (SC), a sensorimotor midbrain structure, plays an important role in sound localization and has been shown to have a topographic map of the auditory space in a range of species. In mice, previous studies using broadband white noise stimuli found that neurons use high-frequency monaural spectral cues and interaural level differences (ILDs) to compute spatially restricted receptive fields (RFs), and that these RFs are organized topographically along the azimuth. However, in a naturalistic environment, the auditory stimuli that an animal encounters may have rich spectral components; however, these sound sources can still be localized efficiently. It remains unknown whether and how the SC neurons respond to naturalistic sounds and, in turn, compute a spatially restricted RF. Here, we show results from large-scale in vivo physiological recordings of SC neurons in response to white noise, naturalistic ultrasonic pup calls and chirps. We find that mouse SC auditory neurons respond to pup calls with distinct temporal patterns and a spatial preference predominantly at ∼60 degrees in contralateral azimuth. In addition, we categorized auditory SC neurons based on their spectrotemporal receptive field patterns and demonstrated that there are at least 4 distinct subtypes of auditory responsive SC neurons.

**Significance Statement:** The superior colliculus (SC) receives visual and auditory information that is used to localize objects. While the organization and composition of visually responsive SC neurons is well described, much less is known about the types and response properties of auditory SC neurons. Here, we presented white noise, ultrasonic mouse pup calls, and chirp stimuli to mice while recording from SC neurons. Analysis of neuronal responses defines 4 distinct classes of auditory neurons. We also show that while auditory neurons respond to naturalistic stimuli, these responses mainly occur when presented from the side but not the front of the animal. These results lead to the hypothesis that mice use different strategies to localize sound depending on the spectral composition of the source.

## Introduction

The superior colliculus (SC) is a sensorimotor midbrain structure that receives and integrates visual, auditory and somatosensory inputs and initiates motor commands for orienting behaviors. The SC can be divided into two main layers: the superficial SC (sSC), which contains neurons that are exclusively visually responsive, and the deep SC (dSC), which contains multimodal neurons that respond to visual, auditory, and/or somatosensory stimuli. The visual response properties of the mouse sSC neurons have been well characterized using electrophysiology and calcium imaging to identify 24 functional subtypes (Wang et al. 2010; Inayat et al. 2015; Ito, Feldheim, and Litke 2017; De Franceschi and Solomon 2018; Li and Meister 2023). However, these studies have been mostly limited to the sSC visual neurons (Ito and Feldheim 2018; Cang et al. 2018; Basso, Bickford, and Cang 2021; X. Liu et al. 2022; Cang et al. 2024).

Much less is known about the response properties and subtypes of the dSC auditory-responsive neurons. It has been shown that there are three types of neurons in the dSC (spike-adapting, late-spiking and fast-spiking) based on their electrophysiological and morphological properties (Bednárová, Grothe, and Myoga 2018), but because this was done in slice culture, it is not known if these neurons are auditory responsive. Thus far, there have been no comprehensive in vivo studies of auditory SC neurons as has been done for the visually responsive neurons in the sSC.

Recent studies have provided new insights into how the SC auditory neurons use binaural and monaural auditory cues to compute spatially restricted receptive fields (RFs). In mice, about half of the neurons in the dSC respond to sound, and ∼23% of these have spatially restricted RFs (Ito et al. 2020; Si et al. 2022). These neurons are organized as a map of sound location in the dSC, with neurons in the anterior dSC responding to sound from the front of the animal and posterior dSC neurons responding to sound from the contralateral side (Ito et al. 2020). Interestingly, SC neurons with frontal and lateral RFs use different cues to compute their RFs; neurons with frontal RFs use monaural spectral cues, while neurons with lateral RFs use Interaural Level Differences (ILDs). One seeming disadvantage of localizing sound using spectral cues is that the spectra of the sound that arrives at the eardrums are influenced by both the spectra of the source sound and the modulation of the spectra due to the shape of the animals’ ears and head, but the brain cannot distinguish between them. Therefore, if the spectrum of the source sound has rich spectral components, it is hypothesized that spectral cues will be unable to provide accurate information about the sound source location; thus, animals may need to rely on ILDs or to develop other strategies to localize such stimuli. In fact, many forms of naturalistic sounds that convey important information for an animal are frequency-rich, and behaviorally, it has been shown that animals are able to localize such sounds. For example, mothers can localize their pups that are making ultrasonic vocalizations (pup calls) that have a rich set of ultrasonic frequencies (Smith 1976; Smotherman et al. 1974).

To better understand how SC neurons respond to and localize sounds, we designed experiments to characterize the auditory response properties of SC neurons to a variety of auditory stimuli. These stimuli included white noise, naturalistic ultrasonic pup calls, and chirps, and were presented to head-fixed mice while recording the electrophysiological responses of a large population of neurons in the dSC. We characterized the temporal response patterns and found that mouse SC auditory neurons respond to pup calls with distinct temporal patterns, suggesting the existence of distinct functional subtypes of auditory neurons. We then determined the spatial RFs of SC neurons when responding to naturalistic pup calls and found a consistent tuning for spatially restricted pup calls coming from ∼60° contralateral azimuth.

To better understand the functional subtypes of auditory neurons in the SC, we determined the spectral-temporal receptive fields (STRFs) of auditory responsive neurons and showed that the STRFs can be used to categorize SC neurons into at least 4 distinct classes. We also show that the STRF subtype identity can be used to predict how a neuron responds to a variety of auditory stimuli. These results provide new insights into how SC neurons localize sounds and demonstrate a way to identify subtypes of auditory responsive neurons in the SC.

## Results

### SC neurons respond to pup calls and show several temporal response patterns

To determine whether and how SC neurons respond to ultrasonic pup calls, we presented pup call stimuli from different azimuthal locations of virtual auditory space (VAS) to awake adult CBa/CaJ mice that were allowed to locomote on a cylindrical treadmill (Figure 1A, B) and recorded the responses of SC neurons using a four-shank (64 electrodes/shank) silicon probe. We used previously measured HRTFs (see methods, (Ito et al. 2020)) to filter a pre-recorded pup call stimulus (Fig 1B) and presented it with calibrated earphones so that the mouse would hear the sound as if it came from one of 17 azimuths (−144° to 144° with 18° spacing) in the horizontal plane. In recordings from 8 mice (P63 - P111, 4 males and 4 females), we recorded the spiking activity of 1501 neurons in the dSC (300–1600 µm from the surface of the SC) where the bulk of the auditory responsive SC neurons reside (Ito et al. 2020).

**Figure 1:**
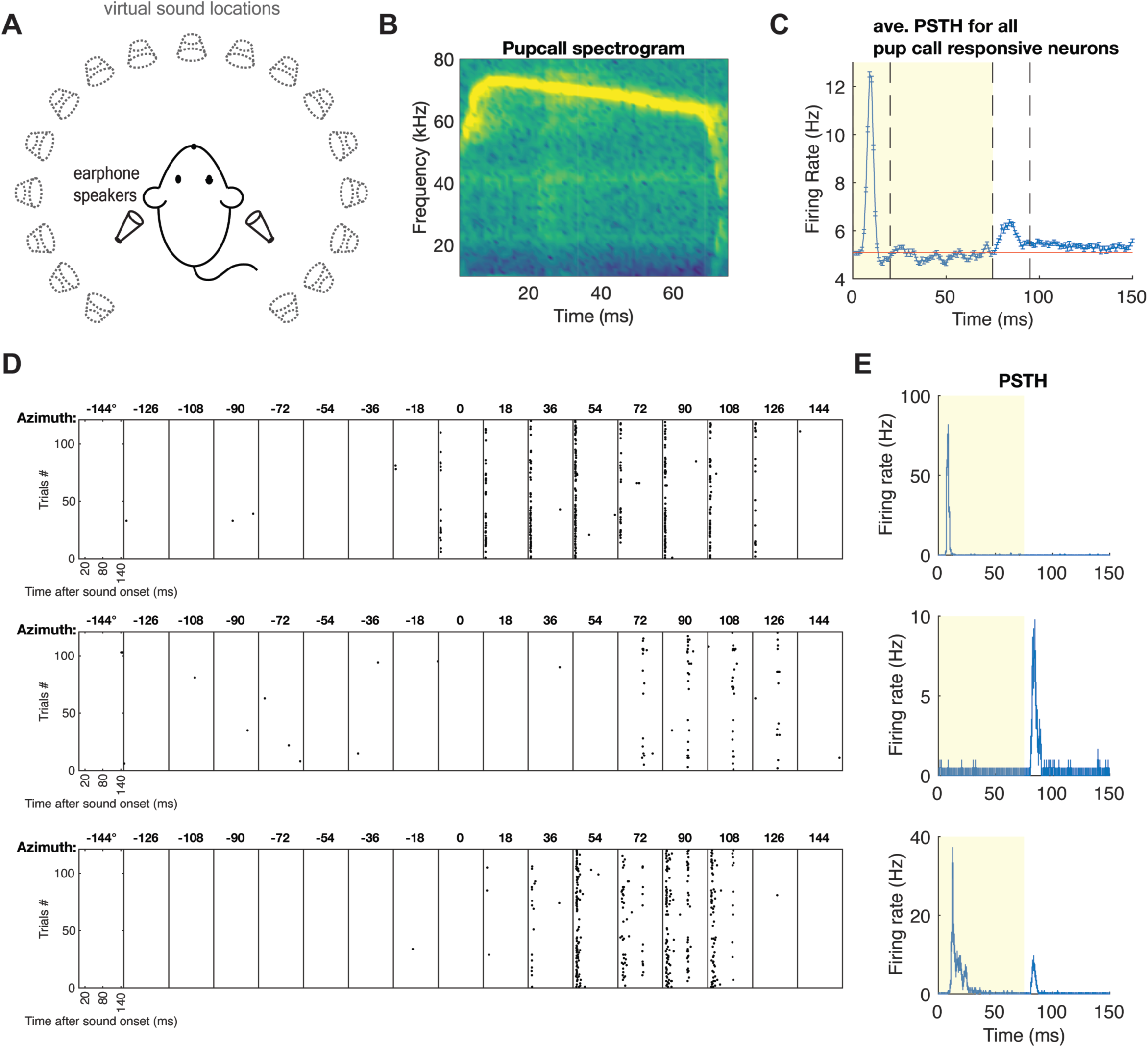
Pup call experiment setup and example responsive neurons in the SC. **A. B** schematic of the experiment setup: head-fixed mice were presented with virtual pup call stimuli at 17 azimuthal locations at 0° elevation with earphone speakers pointing to the ear canals. ***B***. The spectrogram of a typical pup call stimulus that was used in this study. ***C***. The temporal response to the pup call stimulus of SC neurons that are responsive to the pup call stimulus. (The red vertical line marks the end of the stimulus; the dashed gray lines mark the first 20 ms of the onset and offset of the stimulus.) ***D***. Raster plot of three example neurons with onset and/or offset response to localized pup call stimuli. ***E***. Peri-stimulus time histogram (PSTH) of the three examples of pup call responsive neurons in ***D***. (The shaded area represents the presence of the stimulus. This is an average response across all azimuths for each neuron).

We found many neurons with significant responses to the onset and/or offset of the pup call stimulus. Analysis of the overall temporal response pattern reveals that, on average, neurons fire in response to the onset and/or offset of the stimulus within 20 ms. To identify the neurons that are responsive to the pup call stimulus, we compared the sound-evoked firing rate with the baseline firing rate for each neuron at each azimuthal location during the onset (0-20 ms), offset (75-95 ms) and in between (20-75 ms) time windows, and identified neurons that significantly responded to the pup call stimulus from at least one spatial location during at least one time window using Poisson statistics (see Methods). We found (30.2 ± 1.2) % (453) of recorded neurons are pup call responsive, some have only an onset response (Fig 1D-E top), some have only an offset response (Fig 1D-E middle), and some that exhibit both an onset and offset response (Fig 1D-E, bottom).

To further characterize the response properties of pup-call responsive neurons, we performed clustering analysis on the response patterns of the 453 pup-call responsive neurons in our data set. To do this, we first normalized the peristimulus time histogram (PSTH) pattern for each pup call responsive neuron (Fig 2A), then we concatenated the normalized response of all pup call responsive neurons to build the response pattern matrix (Fig 2B) for further clustering analysis using principal component analysis (PCA) and k-means clustering (see methods). This resulted in five clusters in the principal component (PC) space (Fig 2C, shown in the first 3 PC axes), and sorted and labeled in the response pattern matrix (Fig 2D). Within each cluster, the neurons show similar temporal response patterns; therefore, we named these clusters based on the latency of the peak of their response: no peak (neurons that do not have an obvious peak), fast peak (neurons that have an onset peak around 10 ms), mid peak (neurons that have a peak in the middle, between onset and offset response), slow peak (neurons that have an onset peak 10 ms or slightly later), and late peak (neurons that have an offset peak around 85 ms).

**Figure 2:**
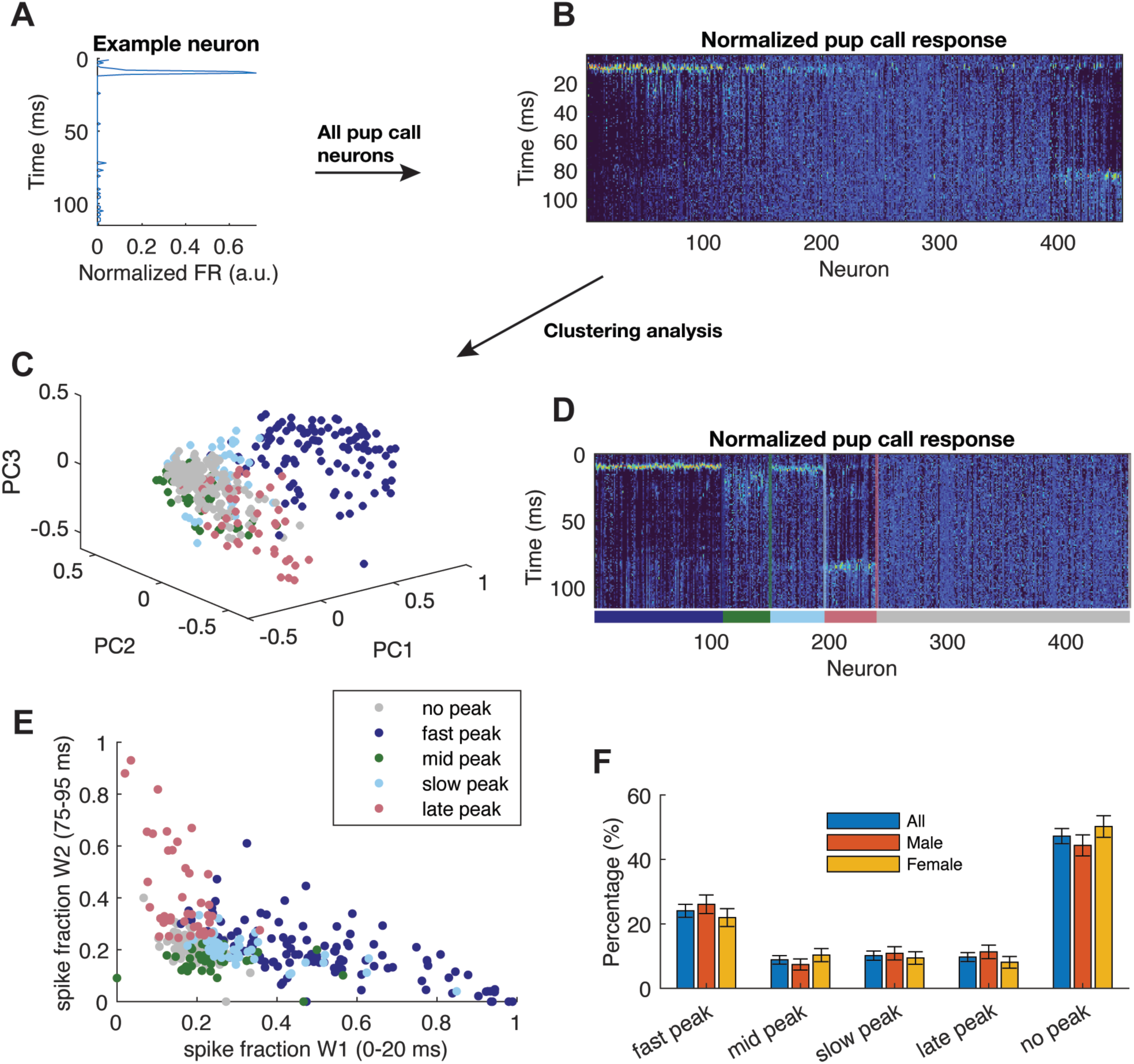
SC neurons respond to pup call stimulus with various temporal patterns. ***A-B.*** Construction of the normalized pup call response matrix for all pup call responsive neurons. Each pup call responsive neuron’s PSTH is normalized (***A***) and concatenated with all other pup call responsive neurons to construct a matrix which is then used for clustering analysis. (Neurons are sorted by the center of mass of their PSTH in ***B***.) ***C***. Results of clustering analysis shown with the first three principal component axes. The clusters are color coded as follows: dark blue - fast peak, green - mid peak, cyan - slow peak, pink - late peak, gray - no peak. ***D***. The normalized pup call response matrix sorted by the results from the clustering analysis, in which the five identified clusters are color coded as in ***C***. ***E***. The same clustering results in ***C*** can be characterized with the fraction of spikes within 0-20 ms vs 75-95 ms for each neuron. ***F***. There’s no difference between sex in terms of the percentage of neurons in each cluster.

To verify the temporal properties of these clusters, we then measured the number of spikes for each neuron within the onset (0-20 ms) and offset (75-95 ms) response windows and calculated the fraction of spikes from each neuron within the two time windows (Fig 2E). Indeed, the fast/slow peak and late peak neurons stand out as having a higher fraction of spikes in one of the on/offset windows. Neurons from the same cluster are also localized close to each other in this plot, indicating that the clustering results reflect the similarity in the peak time latencies for the pup call responsive neurons.

Pup calls are known to trigger maternal behaviors in mice, and previous studies have shown differences in the behavioral response to pup calls between male and female mice (Carcea et al. 2021; Marlin et al. 2015). To determine if the SC neuronal response to pup calls varies between sexes, we compared the fractions of neurons that belong to each of the clusters. We found that these classes of temporal response properties were present in both male and female mice with no significant difference in the percentage of each type (Fig. 2F).

### Neurons that are responsive to both white noise and pup call stimuli have a topographic organization with white noise but not pup call stimulation

We next wanted to determine if pup-call responsive neurons respond exclusively to pup calls (Fig 3A-D). Therefore, we examined the responsiveness of SC neurons to white noise and pup call stimuli (Fig 3D, see methods). We find that (70 ± 1.2 )% of all the recorded SC neurons respond to either white noise or pup call stimulus; more than half of these ((57 ± 2)%) respond to only white noise, about a third ((33.4 ± 1.7)%) respond to both the white noise and pup call stimuli, and only a small fraction ((9.8 ± 0.9)%) respond to the pup call but not white noise stimulus.

**Figure 3:**
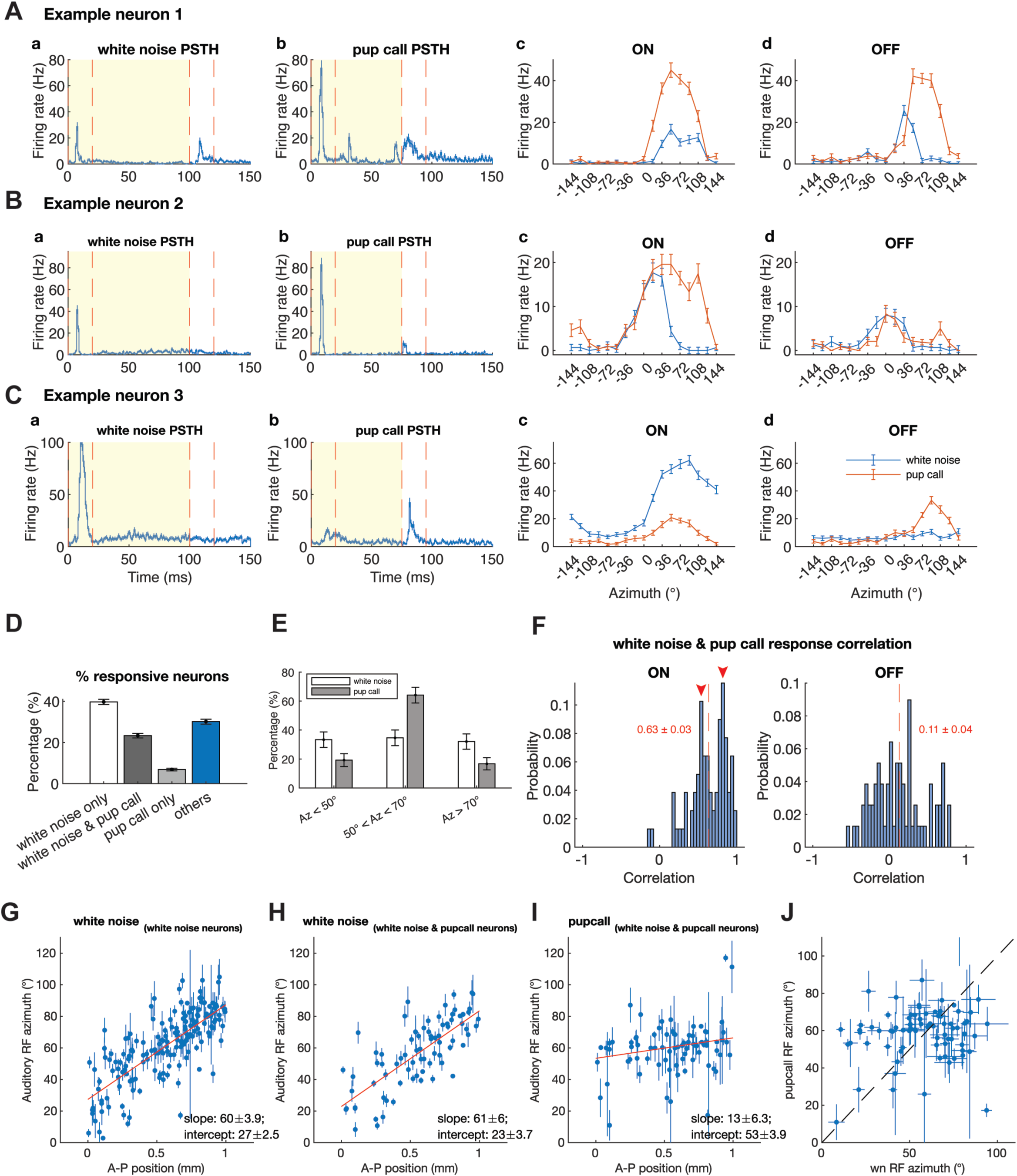
The topographic organization of pup call and white-noise responsive neurons in the SC. ***A-C.*** Example neurons that are responsive to both white noise and pup call stimuli. PSTHs for their white noise response (***a***) and pup call response (***b***), and azimuthal RFs of their onset (***c***) and offset (***d***) response to white noise (blue) and pup call (red) stimuli. ***D.*** Percentage of neurons that respond to white noise and/or pup call stimuli, out of all neurons recorded. ***E.*** Percentage of neurons that have a RF azimuth centered at less than 50°, between 50° and 80°, or more than 80° when responding to white noise (white) and pup call (gray) stimuli, out of all neurons that significantly respond to both. ***F.*** Histograms of the correlation between the azimuthal RFs of white noise and pup call stimuli for neurons that are responsive to both in their onset (left) and offset (right) response. (The red dash line marks the mean, and the red numbers are (mean ± SEM) of the distribution. Red arrowheads in the left panel mark the two peaks in the distribution) ***G***. The topographic map of the auditory space from neurons that are significantly responsive to white-noise stimuli, measured with localized white-noise stimuli. ***H***. The topographic map of the auditory space from neurons that are significantly responsive to both white-noise and pup call stimuli, measured with localized white-noise stimuli. ***I***. The RF centers measured with localized pup call stimuli as a function of the anteroposterior (AP) SC positions of neurons significantly responsive to both white-noise and pup call stimuli. ***J***. The center of spatially restricted RFs of neurons that are significantly responsive to both localized white-noise and pup call stimuli, tested with localized white-noise and pup call stimuli. (error bars: fitting error from the Kent (***G***) or Gaussian (***H-J***) distributions; red lines in ***G-I***: linear fitting results, slopes and intercepts of the lines shown in the plot; the dashed line in ***J*** marks when the centers of RFs overlap perfectly).

We then determined the spatial RFs along the azimuthal axis of the neurons that responded to both the white noise and pup call stimuli. To measure the azimuthal RFs of neurons that respond to the white noise or pup call stimulus, we calculated the mean firing rate within the onset (0 - 20 ms) and offset (75 - 95 ms) time windows across all trials at each azimuthal location (Fig 3A-C, c-d). We observed RF location matching white noise and pup call azimuthal RFs in neurons with an onset response (Fig 3A, c) and in neurons with an offset response (Fig 3B, d). We also observed that some neurons had RF locations that were shifted along the azimuth when responding to white noise versus the pup call stimuli. For example, neurons like example neuron 2 (Fig 3B,c) responds to the onset of the white noise stimulus from a much more limited range of azimuths restricted to the front of the mouse (Fig 3B, c), but its pup call azimuthal RF is much broader and centered more laterally. Alternatively, neurons like example neuron 3 respond to the onset of the white noise stimulus from a much broader range of azimuths on the contralateral hemisphere while the pup call onset response still favors the ∼60 degree contralateral azimuth (Fig 3C, c), resulting in a pup call azimuthal RF center shifted more laterally. To quantify how much pup call azimuthal RFs match with the white noise RFs for SC neurons, we calculated the correlation between the firing rate at the 17 azimuthal locations when responding to the white noise versus the pup call stimulus (Pearson correlation, Fig 3F shows the probability distribution of the correlation coefficient). We found that the pup call and white noise azimuthal RFs have a much higher correlation in the onset response than in the offset response. Interestingly, we observed two peaks in the probability distribution of the correlation coefficient measured from the correlation between the white noise and pup call azimuthal RFs (Fig 3F, left, red arrowheads). This indicates the existence of potentially two populations of neurons: a group of neurons with matching white noise and pup call azimuthal RFs (Fig 3F, left, the right arrowhead, centered ∼0.8), and another group of neurons with shifted white noise and pup call azimuthal RFs (Fig 3F, left, the right arrowhead centered ∼0.6).

We previously proposed that neurons in the anterior SC compute frontal RFs using the ∼25-80 kHz range of the monaural spectral cues while neurons in the posterior SC compute lateral RFs using ILDs. Because pup calls do not contain the full 40-60 kHz spectrum (Fig 1B) we hypothesized that frontal space would not be represented by pup call stimulated neurons’ RFs and, therefore, there would not be a topographic map of sound space of pup call responses. To test this hypothesis, we graphed the location of each pup call neuron’s spatial RF vs. its A–P location, determined the slope of the best fit line and compared this to a similar graph generated by the white noise response. Consistent with our previous studies, we find that the spatial RFs of neurons using the white noise stimuli are organized topographically along the A-P axis of the SC (Fig. 3G) with a slope of (60 ± 3.9)°/mm and intercept of (27 ± 2.5)° (Ito et al. 2020, 2021; Si et al. 2022). This does not significantly change if we limit our analysis to the subset of neurons that responded to both white noise and pup call stimuli (Fig 3H) (slope: (61 ± 6)°/mm, intercept: (23 ± 3.7)°). However, when we plot the azimuthal RF centers against the A-P positions of the same neurons responding to a pup call stimulus, we find that the slope of the line is (13 ± 6.3)°/mm, significantly less than that determined using a white noise stimulus. (p = 9x10^-11^, ANOVA). Instead of having a distribution of RFs from the most frontal to the back of the azimuthal axis as the white-noise topographic map, we observed that a majority of the pup call RFs are centered at ∼60° azimuth (Fig 3J) with very few monitoring sound coming from the front ((19 ± 4)% compared to (33 ± 5)% for white noise) or the more lateral space ((17 ± 4)% compared to (32 ± 5)% for white noise) (Fig 3E). Therefore, although the dSC neurons can “hear” pup calls, they likely interpret them to be coming from contralateral auditory space regardless of the true location of the sound.

### There are at least 4 classes of SC auditory neurons based on responses to random chord stimuli

We show that there are five classes of auditory SC neurons based on their temporal response patterns to pup call stimuli (Figure 2). In addition, we demonstrate that the same neuron can respond differently to white noise and pup call stimuli (Figure 3). Because pup calls differ from white noise stimuli in their spectrotemporal structures, this suggests that the SC neurons may process spectrotemporal information in distinct ways. This information can be used to determine functional subtypes of the auditory SC neurons.

To classify neurons based on their spectrotemporal responses, we used random chord stimuli to determine each neuron’s spectrotemporal receptive field (STRF, see methods). To do so, we recorded the spiking response of neurons while the animals were presented with dynamic random chord stimuli independently to each ear using frequencies ranging from 5 to 80 kHz with each random chord lasting 5 ms (see methods). We then determined the spike triggered averages (STAs) from stimuli presented at each ear. By comparing the STAs with the average probability of the presence of each given tone, we determined which pixel(s) in the 48 (number of tones) by 10 (number of time bins) by 2 (contra- or ipsi-lateral sides) STA images are significantly more positive or negative than chance alone. Consistent with our previous findings (Ito et al. 2020; Si et al. 2022), we find the majority of both positive (Fig 4A, left) and negative (Fig 4A, right) significant pixels are > 20 kHz. There is also a contralateral bias for both the positive and negative significant pixels (Fig 4A and 4C left). Numbers and positions of significant STRF pixels for individual SC neurons are sparse and highly variable, which can be inferred from the exponential decrease in the number of pixels when progressively increasing the threshold of the minimum number of significantly responsive neurons (Fig 4B). Out of all neurons recorded, we found 12.7% (190 out of 1501) neurons that have at least one significant STRF pixel, which are therefore considered to have a significant STRF.

**Figure 4:**
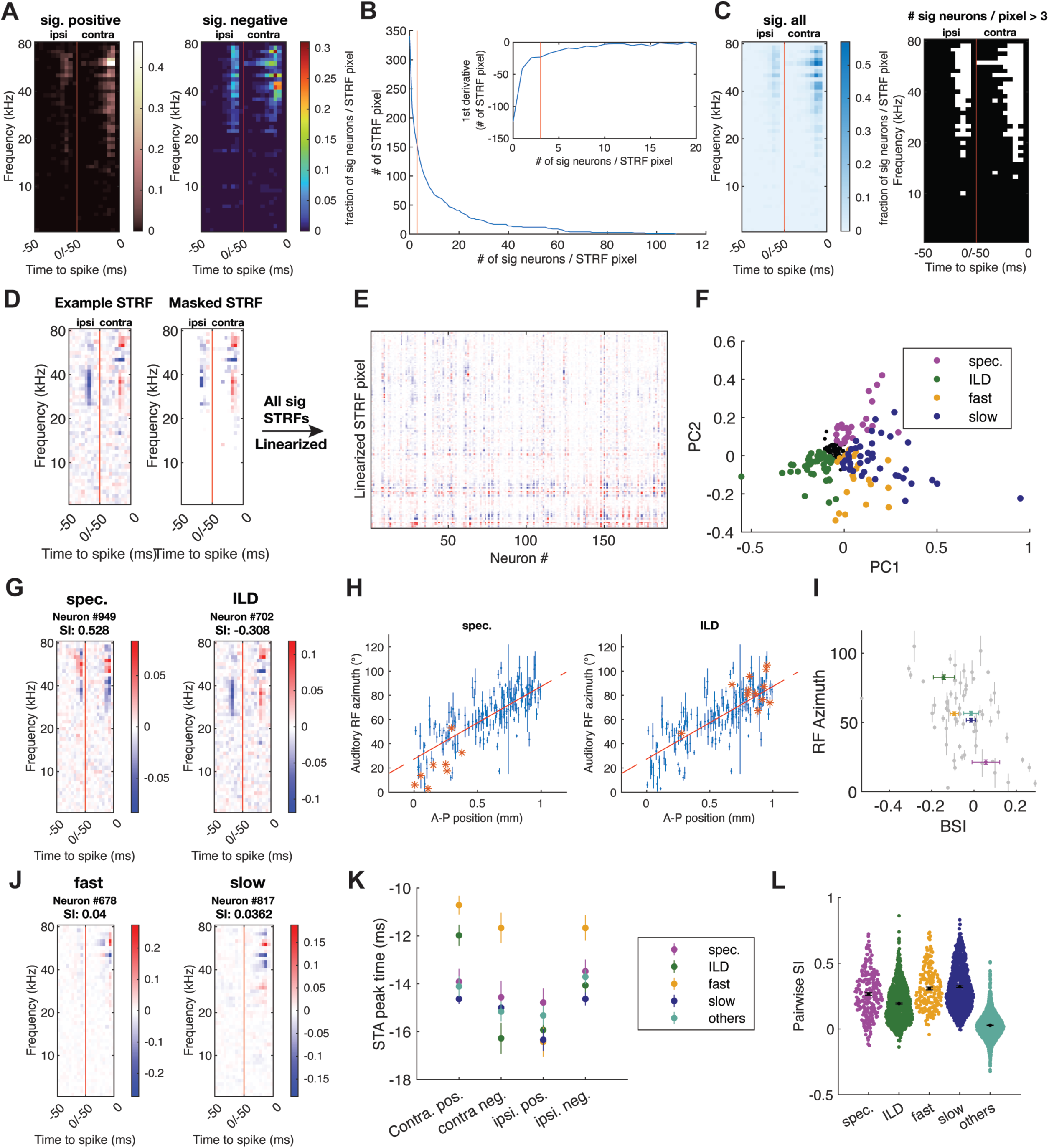
Clustering analysis of STRF patterns reveals at least four subtypes of auditory SC neurons. ***A-F***. Clustering analysis of STRFs of SC neurons. ***A.*** Heatmap showing the probability of SC neurons showing significant positive (left) or negative (right) response at each spectral-temporal pixel in their STRFs. The red line marks the border between contralateral and ipsilateral sides. ***B.*** Number of STRF pixels, whereby the SC neurons have a significant response, as a function of the number of significantly responsive neurons per pixel. Inset, the first derivative of the curve; red vertical line, the threshold used in the following clustering analysis (# of sig neurons / STRF pixel = 3). ***C.*** Left: Heatmap showing the probability of SC neurons with significant response (positive and negative combined) at each spectral-temporal pixel in their STRFs. The red line marks the border between contralateral and ipsilateral sides. Right: The mask used for the following clustering analysis. White, pixels where there are more than 3 neurons significantly responsive. The red line marks the border between contralateral and ipsilateral sides. ***D.*** An example STRF before and after the application of the mask in ***C***. ***E.*** The matrix used for clustering analysis consists of linearized and masked STRF pixels from all neurons that are significantly responsive to the random chord stimulus. ***F***. Results of clustering analysis shown with the first two principal components. ***G.*** STRFs of two example neurons from the clusters of spectral cue neurons (spec.) and ILD neurons (ILD). ***H***. RF azimuth and AP position of neurons from the spec. and ILD clusters relative to all neurons in the topographic map of the auditory space. (red asterisks: highlighted neurons in the spec. (left) and ILD (right) clusters that also significantly respond to localized white-noise stimuli; blue dots, error bars and red lines: all neurons that significantly respond to localized white-noise stimuli with errors from the Kent distribution fitting and linear fitting results as in ***Fig 3I***). ***I***. Average RF azimuth and average similarity index (SI) for neurons from each STRF cluster (gray points are all neurons that have significant STRF and localized RFs, error bars are SEM of neurons from each cluster; color code as in ***F***). ***J.*** STRFs of two example neurons from the clusters of fast neurons (fast) and slow neurons (slow). ***K.*** average peak time in the STRF for positive or negative response on the contralateral or the ipsilateral side for the four STRF clusters: spec., ILD, fast and slow. L. The distributions of pairwise SI values for each possible pair of neurons within the four STRF clusters. Black dots and error bars, mean ± SEM. spec., the spectral cue neurons from the STRF clustering analysis; ILD, neurons from the ILD STRF cluster; fast, neurons from the fast STRF cluster; slow, neurons from the slow STRF cluster.

To identify functional subtypes of SC neurons, we then did cluster analysis on the STRF patterns. We first created a mask to only select and linearize only the most significant STRF pixels. We used the threshold of including only pixels with more than 3 neurons being significant (Fig 4B-D), which resulted in 153 pixels for each neuron. We then constructed a matrix with the linearized 153 pixels from all neurons that have a significant STRF (Fig 4E). Clustering analysis revealed four distinct classes of neurons (Fig 4F): neurons that have highly similar contra/ipsi-lateral STRF patterns (spectral cue neurons, example STRF shown in Fig 4G, left); neurons that have anti-correlated contra/ipsi-lateral STRF patterns (ILD neurons, example STRF shown in Fig 4G, right ); neurons that have transient response to sound (fast, example STRF shown in Fig 4J, left); and neurons that have relatively prolonged response to sound (slow, example STRF shown in Fig 4J, right). There is a fifth class of neurons (others, black points in Fig 4F and cyan points in Figs I and L), but when we calculated the pairwise similarity index (pairwise SI, Fig 4L) to measure the similarity of STRFs from the same class of neurons, we found a significantly low similarity between neurons that belong to that class of neurons. This indicates that these neurons have a mixture of different STRF patterns; therefore, we have excluded this class in further analysis.

### The Spectral Cue and ILD classes defined by STRF analysis are distinct functional subtypes

If the four classes of auditory neurons identified using STRFs are functional subtypes we would expect that neurons within each class would share other properties such as their location in the SC, and their responses to other stimuli. When we determined the physical location of each class of neurons in the SC, we find that neurons in the spectral cue class are predominantly located in the anterior SC and have frontal spatial RFs (Fig 4H, left), while neurons in the ILD class are predominantly located in the posterior SC and have lateral spatial RFs (Fig 4H, right). The fast and slow classes were distributed throughout the A-P axis of the SC and have RF azimuth RFs that lie between the spectral cue and ILD classes.

We examined all neurons that have both a significant spatially restricted RF and a significant STRF and calculated each neuron’s binaural cosine similarity index (BSI, see methods), which reflects how similar their STRFs are between the contralateral and ipsilateral side. Consistent with our previous work (Ito et al. 2020), the RF azimuth and BSI are anti-correlated (Fig 4I). Further analysis reveals that spectral cue and ILD neurons belong to different classes, the average value for RF azimuth and BSI of these two classes of neurons are well separated in the RF azimuth over BSI plot (Fig 4I, green and purple data points and error bars). Notably, neurons from the fast or slow classes have BSI and RF azimuth values in between the spectral cue and ILD classes, but these two classes of neurons cannot be well separated based on their A-P position and their RF azimuths (Fig 4I). Instead, neurons from the fast class stand out as having the shortest latency to peak time in their STRFs (Fig 4K). We measured each neuron’s latency to peak time for both the contralateral and ipsilateral sides of their STRFs by taking the average of both the positive and negative responses across frequencies and determining the peak time for each averaged response. We found that neurons from the fast class are distinguished by their significantly faster peak time in both positive and negative responses on the contralateral side, and the negative response on the ipsilateral side (Fig 4K, yellow data points and error bars).

We also determined the response properties of the classified neurons to white noise (Fig 5A, a) and non-broadband stimuli such as pup calls (Fig 5A, b), and chirps (20-80 kHz of different frequency/time slopes (Fig5A, c-f). Figure 5A shows the average PSTH of all neurons and Figure 5B organizes the data such that the neurons in each group defined by their random chord response (spectral cue, ILD, fast, slow), and that also responded to the other stimuli, are grouped together (white lines). Inspection of the data shows that while virtually all neurons fire in response to the onset of the stimulus, very few neurons fire in response to the offset of a stimulus, except for neurons in the ILD and to some extent the fast class (Fig 5B, D). For example, in response to the white noise stimulus (5Aa-Ea) all neurons have a fast onset response, which is similar to neurons in the four STRF classes but have very little response to the offset of the stimulus. After presenting a pup call stimulus, we see the fast onset response that is present for most neurons from the four STRF classes; however, neurons from the ILD class have more of an offset response than the other classes (Fig 5B, b, second block from the left in the heatmap). To quantify this, we calculated the following for each STRF class: 1) the percentage of neurons that significantly respond to the offset of the stimulus (Fig 5D, a-f), and 2) for each neuron within each STRF class, the offset response amplitude measured by the z-score within 20 ms after the offset of the stimulus (Fig 5E, a-f). We find that there is a significantly higher percentage of ILD neurons that respond to the offset of all frequency rich stimuli (but not white noise), and that the ILD neurons have a higher response amplitude when responding to the offset of these stimuli (Fig 5D&E, b, p < 1x10^-5^ for both). Indeed, when we mark neurons from the four STRF classes in the first two principal component (PC) space of white noise and pup call response, the neurons from the four STRF classes are intermingled in the white noise PC space (Fig 5C, a), but are more separated in the pup call PC space (Fig 5C, b). This suggests that the response patterns of SC neurons to pup call, but not white noise, can be classified by their STRF patterns.

**Figure 5:**
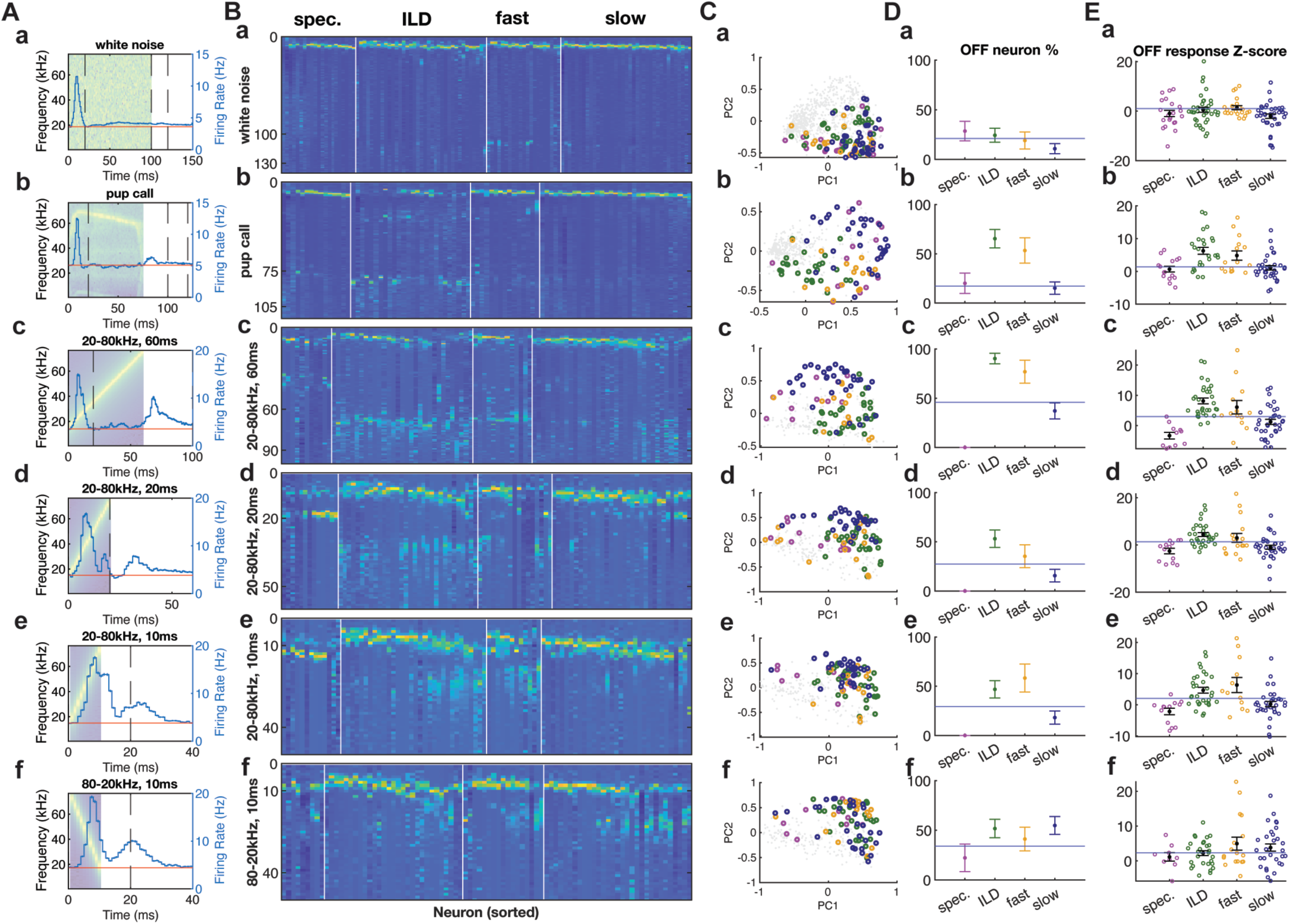
The temporal response patterns of SC neurons to white noise, pup call and chirp stimuli separated by the STRF labels. ***A.*** The PSTH of neurons responding to white noise (***a***), pup call (***b***), 60 ms chirp that goes from 20 to 80 kHz (20-80 kHz 60ms, ***c***), 20 ms chirp that goes from 20 to 80 kHz (20-80 kHz 20 ms, ***d***), 10 ms chirp that goes from 20 to 80 kHz (20-80 kHz 10 ms, ***e***), 10 ms chirp that goes from 80 to 20 kHz (80-20 kHz 10 ms, ***f***) with the spectrogram of each stimulus in the background. ***B***. Temporal response of neurons in the four identified STRF clusters (spec., ILD, fast, slow, separated by the white lines) to variations of pup call and chirp stimuli as in A. (y axis, time/ms). ***C***. Neurons from the four identified STRF clusters (spec., ILD, fast, slow) highlighted on the first two principal component axes of all neurons significantly responsive to both dynamic random chord and at least one of the tested variations of pup call and chirp stimuli. (Black dots: all significant neurons; color code as in Fig 4.) ***D***. Percentage of neurons in the four STRF clusters that exhibit significant offset response. (y axis, percentage/%; horizontal line, average percentage of neurons that show offset response out of all neurons that are significantly responsive to each stimulus tested.) ***E***. The offset response amplitude shown in Z-score for neurons in the four STRF clusters. (y axis, Z-score in 20 ms bin; horizontal line, average offset response amplitude of neurons that are significantly responsive to each stimulus tested; black points and error bars, mean ± SEM of each cluster; spec., the spectral cue neurons from the STRF clustering analysis; ILD, neurons from the ILD STRF cluster; fast, neurons from the fast STRF cluster; slow, neurons from the slow STRF cluster.)

To further test the hypothesis that non-broadband stimuli elicit different response patterns from neurons of distinct STRF classes, we extended our stimuli and collected data from SC neurons responding to four types of chirps with frequencies between 20 and 80 kHz but with different frequency/time slopes. The average PSTHs of SC neurons responding to these chirps show consistent onset and offset responses within the 20 ms after both onset and offset of the stimuli (Fig 5A, c-f). We then obtained the sorted normalized response matrix for each of these chirp stimuli, as we did for the white noise and pup call response, such that neurons from each STRF class are grouped together (Fig 5B, c-f). The responses of neurons to these stimuli are more similar to those from the pup call stimulus than the white noise stimulus, namely, neurons from different STRF classes tend to have distinguishably different temporal response patterns.

This can be seen when we display the neurons from the four STRF classes in the first two PC spaces for each of the chirp responses (Fig 5C, c-f). Neurons from different STRF classes tend to be clustered instead of being intermingled as in the white noise response. Specifically, neurons from the ILD class also stand out as having more offset responses in three of the four chirps (the exception being a chirp from 80 kHz to 20 kHz with a duration of 10 ms). Indeed, the percentage of offset responsive ILD neurons and their response amplitude are significantly higher than average (Fig 5D, c&d, p < 1x10^-5^, e, p = 0.0051; Fig 5E, c&d, p < 1x10^-5^, e, p = 0.0004). In addition, we consistently found more offset responsive neurons in the fast STRF class. Finally, we see a strikingly low percentage of offset responsive spectral cue neurons (Fig 5D, c-e, p < 1x10^-5^ for all), and their offset response amplitude is also lower than average (Fig 5E, c, p < 1x10^-5^, d, p = 0.0009, e, p = 0.0001) for all stimuli tested. Taken together, these results show that the spectral cue and ILD classes of neurons identified solely by using STRFs represent distinct functional classes.

## Discussion

In this study, we show that mouse SC auditory neurons respond to pup calls with distinct temporal patterns and a spatial preference predominantly at ∼60 degrees in contralateral azimuth. We also demonstrate a way to classify SC neurons into at least 4 distinct subtypes using the STRF patterns.

Previous studies have shown that response to pup calls in the primary auditory cortex is enhanced only in female mice with maternal experience (Carcea et al. 2021; Marlin et al. 2015). However, we did not find differences between sexes in the temporal response patterns of SC neurons that responded to pup calls. This suggests that pup calls are processed rather similarly in the SC auditory neurons of virgin female and male mice, consistent with the role of the SC in determining the location but not the identity of an auditory stimulus. One limitation of our experiments is that we only used virgin mice in this study. Given the evidence that mother mice or experienced female virgin mice exhibit different interactive behavior with pups (Carcea et al. 2021), it will be interesting to examine the differences between neuronal responses to pup calls in the SC of experienced mothers and virgin females. Another limitation of this finding is that we only compared the differences between sexes in terms of the temporal response patterns of SC neurons. It is possible that differences exist in other aspects that we did not characterize.

We did not observe a topographic map of the auditory space measured with pup calls in the SC, even though the same population of neurons produced a topographic map when measured with the white noise stimulus. We showed that neurons with white noise RFs centered at the most frontal or lateral auditory space have mismatching pup call RFs: the frontal white noise RFs shift laterally when the neurons respond to pup calls, while the lateral white noise RFs shift frontally. Indeed, the probability distribution of the correlation coefficient between the white noise and pup call spatial RFs captures two neuron populations: one with more correlated white noise and pup call RF azimuths, and the other with less correlated RFs (Fig 3F, left). We find that SC auditory neurons respond to pup calls with a spatial preference predominately at ∼60 degrees in contralateral azimuth (Fig 3I-J), which is close to the direction where ILD is maximized (Ito et al. 2020). This suggests that mouse SC neurons mainly use ILDs to localize pup calls.

We have previously shown that high-frequency spectral cues are necessary to compute frontal RFs when SC auditory neurons respond to white noise stimuli (Ito et al. 2020; Si et al. 2022). The pup call stimulus we used in this present study, which is also representative for pup calls in general, is a high frequency stimulus but does not consist of the complete high-frequency range (40-80 kHz) needed to generate within ear contrast that is required to compute frontal RFs. The lack of frontal RFs for pup call responsive neurons indicate that the mere presence of a range of high-frequency sound is not sufficient for SC neurons to compute a topographic map of the auditory space. We speculate that the restricted frequency range in a pup call makes it difficult for the brain to analyze the spectra and separate the modulation by HRTF from the spectra structure of the sound source. Therefore, when spectral cues cannot provide accurate information about the sound source location, the SC neurons rely on ILDs to compute their spatial RF.

What does it mean to a mouse when, instead of having a smooth topographic map of azimuthal space as with broadband noise stimuli, most SC auditory neurons are tuned to respond to pup calls primarily at ∼60 degrees in azimuth? Given the behavioral evidence that mother mice are able to localize the sound source of pup calls (G. Ehret 1987; Günter Ehret 2005), the absence of a smooth topographic map of the auditory space does not equate to the inability to localize pup calls; this suggests that different strategies may be used for sound localization depending on its source. Studies done with human subjects showed that the relative movement of the listener and the sound source can resolve the ambiguity in sound localization (Perrett and Noble 1997; Wightman and Kistler 1999). Taken together with evidence from bats (Aytekin, Moss, and Simon 2008) and mice (Mai et al. 2024), this data suggests that sound localization is a sensorimotor process that requires the auditory system to coordinate with the motor system to localize the sound source using dynamic auditory cues.

We identified four main classes of auditory SC neurons based on their binaural STRFs. This clustering analysis recapitulated our hypothesis that the neurons in the anterior SC mainly use spectral cues to compute frontal RFs, whereas neurons in the posterior SC use ILDs to create lateral RFs; neurons in between combine spectral cues and ILDs to create RFs that span between the frontal and lateral auditory space (Ito et al. 2020). Indeed, we found that neurons in the anterior SC have the most symmetric STRFs between ears and frontal RFs, whereas neurons in the posterior SC have the most across-ear contrast in their STRFs and lateral RFs. In between these two distinct classes of neurons, we found two other subtypes of neurons: the “fast” neurons that are transiently activated by auditory stimulation, and the “slow” neurons that can be activated within a prolonged time window on the contralateral side. Although much work has been done to characterize subtypes of visually responsive neurons in the SC morphologically, genetically, and physiologically (Gale and Murphy 2014, 2016, 2018; Y. Liu et al. 2023; Li and Meister 2023), this is the first effort to characterize the subtypes of in vivo auditory response of the mouse SC neurons.

We observed an offset response in SC auditory neurons when they respond to frequency rich stimuli such as the pup call and chirp stimuli. This is the first time that offset response to sound is reported in the SC. However, offset responses to auditory stimuli have been reported throughout the auditory brain across species (Kopp-Scheinpflug, Sinclair, and Linden 2018), most peripherally from the cochlear nucleus (Suga 1964; Young and Brownell 1976; Ding, Benson, and Voigt 1999), to the superior olivary complex (Dehmel et al. 2002; Kulesza, Spirou, and Berrebi 2003), the inferior colliculus (IC) (Kasai, Ono, and Ohmori 2012; Akimov, Egorova, and Ehret 2017; Xie, Gittelman, and Pollak 2007; Casseday, Ehrlich, and Covey 1994), the medial geniculate body of the thalamus (He 2002; Anderson and Linden 2016), to the auditory cortex (Keller, Kaylegian, and Wehr 2018; Recanzone 2000). Offset auditory response is important for duration discrimination (Casseday, Ehrlich, and Covey 1994) and gap detection (Xu, Fu, and Chen 2014) in a more energy efficient way than encoding the entire duration of a sound. We do not know the source of the offset response in the SC, but because the SC receives direct inputs from the IC and auditory cortex (Edwards et al. 1979; Druga and Syka 1984; Covey, Hall, and Kobler 1987; Jiang, Moore, and King 1997; Doubell et al. 2000; Bednárová, Grothe, and Myoga 2018; Benavidez et al. 2021; Issa, Sekaran, and Llano 2023), it is possible that the offset response to pup calls and chirps in the SC are inherited from the IC, within the SC or from the auditory cortex. Intriguingly, we also found that ILD neurons, but not spectral cue neurons, have an offset response. This observation might indicate that the auditory neuron subtypes get inputs from distinct brain regions. In addition, we also observed spatial tuning of the offset response in SC neurons, both to white noise and pup calls.

Intriguingly, the spatial tuning of the offset response to white noise and pup calls is much less correlated than that of the onset response of SC neurons (Fig 3F). This may imply that the onset and offset responses in the SC are inherited from different upstream sources, or that the spatial auditory RFs are refined by the offset of the stimulus, which can be an interesting future direction of research.

## Methods

### Ethics statement

All procedures were performed in accordance with the University of California, Santa Cruz (UCSC) Institutional Animal Care and Use Committee.

### Procedures

We used the mouse head-related transfer functions (HRTFs) to present virtual auditory space (VAS) stimuli and recorded the activity of the collicular neurons from head-fixed, alert mice using 256-electrode multishank silicon probes. Our experimental procedures were described previously (Shanks et al., 2016, Ito et al. 2017, Ito et al. 2020) and are detailed below.

## Auditory stimulation

### HRTF measurement and VAS stimulation

The HRTFs of CBA/CaJ (Ito et al. 2020) mice were measured and the VAS stimuli were generated as previously described (Ito et al. 2020). The measured HRTFs for CBA/CaJ mice and the explanation for their use are available in the figshare website with the identifiers https://doi.org/10.6084/m9.figshare.11690577 and https://doi.org/10.6084/m9.figshare.11691615.

VAS stimuli were generated as previously described (Ito et al., 2020). In brief, each stimulus sound was filtered by a zero-phase inverse filter of the ES1 speaker (Tucker-Davis Technologies), the eHRTF, and the measured HRIR. The VAS stimulus is then delivered pointing toward the animal’s ear canals. We tried to reproduce the physical configurations used for the HRTF measurements so that the acoustic effect induced by this setup is canceled by an inverse filter of the eHRTFs.

### Full-field white noise stimulus

The baseline stimulus pattern was 50dB SPL, 100 ms white noise containing a flat 5-80kHZ spectrum with linear tapering windows in the first and last 5 ms. After applying the filters mentioned above, the full-field white noise stimulus contains grid points of five elevations (0-80° with 20° steps) and 17 azimuths (-144° to 144° with 18° steps), totaling 85 points in the two-directional field. The stimulus was presented every 2s and repeated 30 times per direction at the intensity of 50 dB SPL.

### Random chord stimulus

To measure spectral tuning properties of the neurons with localized RFs, we used dynamic random chord stimuli and calculated the STA of the stimuli for each neuron. The stimuli consist of 48 tones ranging from 5 to 80 kHz (12 tones per octave). Each pattern was either 10 ms or 20 ms long with 3 ms linear tapering at the beginning and the end. In each pattern, the tones were randomly set to either ON or OFF with a probability of 0.5. The total number of tones per pattern was not fixed. The amplitude of each tone was fixed and set to be 50 dB SPL when averaged over time. One presentation of the stimuli was 2-min long and this presentation with the same set of patterns was repeated 20 times to produce a 40-min-long stimulus. Tones from the left and right speakers were not correlated with each other to measure the tuning to contrast between the ears. We did not use a specific HRTF for this experiment, but simply canceled the eHRTF so that the stimulus sound was filtered to have a flat frequency response near the eardrums.

### Pup call stimulus

To get a representative audio sample of a mouse pup call, we recorded from P5-P9 CBA/CaJ mouse pups with a high-frequency bat detector (Echo Meter Touch, Wildlife Acoustics), and selected an audio clip with the most typical spectrogram. To generate the baseline stimulus, filters were then applied to the recorded audio to calibrate the effect from the recording device. After applying the HRTF filters described above, the horizontal pup call stimulus contains 17 azimuths (−144° to 144° with 18° steps) on the horizontal plane. The stimulus was presented every 1 s and repeated 120 times per direction at the intensity of 50 dB SPL.

### Chirp stimulus

To assess the response of auditory SC neurons to simplified synthetic stimuli, we generated a set of chirp stimuli that cover the 60 kHz frequency range of 20 to 80 kHz, at frequency rates of 1, 3 and 6 kHz/second, and 80 to 20 kHz at a rate of -60 kHz/second. After applying the HRTF filters described above, the horizontal chirp stimuli contain 5 azimuths (0°, 36°, 54°, 72°, 90°) that cover a quadrant of the horizontal plane. The stimulus was presented every 1 s and repeated 60 times per direction at the intensity of 50 dB SPL.

## Mice

The CBA/CaJ mice used in this study were the offspring of the mice initially obtained from The Jackson Laboratory (stocks #000664) and bred in the UCSC vivarium. We used two-to four-month mice of each sex.

## Electrophysiology

One day before the recording, a custom-made titanium head plate is implanted on the mouse’s skull to fix the mouse head to the surgery rig and recording rig without damaging or touching the ears. On the day of the recording, mice were anesthetized with isoflurane (3% induction, 1.2 – 1.5% maintenance; in 100% oxygen) and a craniotomy was made in the left hemisphere above the SC (approximately 1 mm by 2 mm in an elliptic shape, about 0.6 mm from the lambda suture). After recovering from anesthesia, the mouse was head-fixed onto the recording rig, where it was allowed to locomote on a rotating cylinder. A 256-channel, 4-shank silicon probe was inserted into the SC with the support of a thin layer of 2% low-melting-point agarose (in saline), and a layer of mineral oil was added on top to prevent the brain from drying. The silicon probe was aligned along the A-P axis. The multi-unit visual RFs were measured as the probe recorded from the superficial SC in all four shanks. After this measurement the probe was lowered until most of the channels were within the dSC (approximately just past the strong visual response in the superficial SC). Then we lowered and raised the probe by 100-120 μm to avoid the probe movement effect of dragged tissue during recording. Recordings were started about 20-30 minutes after inserting the probe and performed in an anechoic chamber.

The probes used for our recordings were kindly provided by Prof. Masmanidis at UCLA (Shobe et al. 2015; Du et al. 2011). We use 256-channel, 4-shank silicon probes (the 256A series). Each shank has a two-dimensional active recording area of about 1 mm x 86 µm, and the shank pitch is 400 µm. The voltage signals collected from all the electrodes are amplified and sampled at 20 kHz using an RHD2000 256-channel recording and data acquisition system (Intan Technologies).

## Experimental design and statistical analyses

### Blind analysis

To validate our results and reduce potential false-positive findings, we performed blind analysis (MacCoun and Perlmutter 2015). First, we analyzed the spiking activity data from half of the neurons randomly sampled from each recording. After exploring the datasets and fixing the parameters in our analysis, we performed the same analysis on the other half of the data to test whether the conclusion holds true on the blinded data. All the results reported in this study passed a significance test in both the exploratory and the blinded data unless otherwise stated. Spike-sorting

We used custom-designed software for spike sorting (Litke et al. 2004), which was also used in our previous published work (Ito, Feldheim, and Litke 2017; Ito et al. 2020, 2021; Si et al. 2022). Raw analog signals were high-pass filtered at ∼313 Hz and thresholded for spike detection. The high-pass part was used for single-unit identification after motion artifact removal. Detected spikes were clustered based on the principal component analysis of the spike waveforms on the seed electrode and its surrounding electrodes. A mixture of Gaussians models was used for cluster identification. Detailed demonstration of the silicon probe schematics, examples of raw electrophysiology data, and demonstration of the spike-sorting process can be found in our previous work (Si et al. 2022).

## Estimating AP location of neurons

We estimated the relative position of the silicon probes using the visual RF positions measured in the superficial SC, as described in our previous work (Ito, Feldheim, and Litke 2017; Ito et al. 2020, 2021; Si et al. 2022). In brief, we measured the positions of the visual RFs on each shank in the superficial SC using multiunit activity and extrapolated the visual RFs to find an A-P position where the visual RF azimuth was 0°; then by assuming the most superficial electrode of the silicon probe was at 300 μm in depth and the insertion angle of the probe into the SC was ∼25°, we were able to estimate the A-P positions of the auditory neurons (Ito et al. 2020, 2021; Si et al. 2022). The position of a neuron relative to the probe was determined by a two-dimensional Gaussian fit to its spike amplitude across multiple electrodes (Ito, Feldheim, and Litke 2017; Ito et al. 2020, 2021; Si et al. 2022). We only analyzed neurons with a positive A-P position in order not to include neurons located outside the SC.

## Significance test for the auditory responses

### White noise response

We used quasi-Poisson statistics for significance tests of the auditory responses of individual neurons (Ver Hoef and Boveng 2007), to deal with the overdispersion of the post stimulus firing rate because of factors such as bursting of the neural activity, as described in our previously published work (Ito et al. 2020). The overdispersion parameter was estimated by the variance of the spike count divided by the mean, which should be 1 if the spiking activity of a neuron is Poissonian. To determine the significance of the response, we first estimated an overdispersion parameter and considered a response to be significant if the *p* value of the neuron’s spike count is below 0.001 [*p* = 1 − CDF(N), where CDF is a cumulative distribution function of the quasi-Poisson distribution and N is the spike count of the neuron].

### Pup call and chirp response

We used Poisson statistics for significance tests of the auditory responses of individual neurons to pup calls and chirp responses. We first estimated the time course of the typical response to auditory stimuli (Fig 1C, Fig 3) and found that the typical response of most neurons happens within 20 ms after sound onset/offset. Therefore, we evaluated the spiking activities in three time windows: onset (0-20 ms after sound onset), offset (0-20 ms after sound offset) and in-between (between 20 ms after sound onset and sound offset, in order to capture the atypical response patterns of some neurons). We measured the baseline firing rate and calculated the Z-score within each time window at each azimuth using Poisson error and selected neurons with a greater than 2 Z-score at any azimuthal location as significantly responsive to the stimulus.

### Dynamic random chord response

We calculated the STA from the spiking response to the dynamic random chord stimulus described above and determined whether an individual neuron had a significant STRF. First, the spikes of each neuron are discretized in time bins the same as the stimulus segment size (5 ms) and the average of the stimulus patterns that preceded each spike was calculated. The stimulus was considered as 1 if there is a tone in the frequency and 0 otherwise. The significance tests for the STRFs were performed by examining whether the difference between the mean of the stimulus and the STA is significant. We used 0.001 as a significance threshold and Bonferroni correction for evaluating multiple time bins, frequencies and contra/ipsilateral input. Neurons with <20 spikes during the stimulus were not analyzed.

## Function fit to estimate the azimuth of the spatial RFs of the neurons

### White noise response

We used a maximum-likelihood fit of the Kent distribution (Kent 1982) to estimate the azimuth of an RF for the full-field white noise stimulus (17 azimuthal x 5 elevational locations), as in our previous work (Ito et al., 2020, 2021; Si et al., 2022). At each point of the directional field, the likelihood value was calculated based on quasi-Poisson statistics (Ver Hoef and Boveng 2007). The error of each parameter was estimated from the Hessian matrix of the likelihood function. As in our previous work, the front-Z coordinate system has a discontinuity of the azimuth across the midline that gives a problem in fitting and interpretation of the data near the midline. We avoided this issue of fitting the data near the midline with the front-Z coordinate system (determined by our HTRF measurement) by switching to the top-Z coordinate system and only used neurons with their elevation smaller than 30° for azimuthal topography, to avoid an area where two coordinate systems are largely different. To achieve stable fits, we set the azimuthal range to be from −144° to +144° and set the elevation range to be from 0° to 90°.

### Pup call and chirp response

We used virtual space auditory stimuli at 17 or 5 azimuthal locations on the horizontal plane for the pup calls and chirp stimuli. For the neurons’ response to these stimuli, we used a maximum-likelihood fit of the Gaussian distribution to estimate the center azimuth of an RF. At each point that we presented the stimuli, the likelihood was calculated based on Poisson statistics. In parallel, we also fitted the data with a uniform distribution. We compared the goodness of the fit between the Gaussian distribution and the uniform distribution with Bayesian Information Criterion (BIC) and only included neurons that showed a better fitting result with the Gaussian distribution. For the stimuli for which we only tested 5 locations on the horizontal plane (0°, 36°, 54°, 72°, 90°, see above for details), in order to avoid problems in the fitting process caused by the unequal spacing, we first interpolated the data and estimated the response to a stimulus from 18° azimuth using the response to 0° and 36° azimuths. We then fitted the data with the 6 azimuths. There is a known defect in our HRTF in the 72° azimuth 0° elevation location, so we compensated the response to stimuli from that direction before doing any fitting analysis. After the fitting process, the neurons with a better Gaussian fitting result were determined and the mu parameter of the optimal Gaussian fit was used as the azimuth of an RF, while the error was estimated from the Hessian matrix of the likelihood function.

## Clustering analysis

### Response to pup calls

We first reduced the dimensionality of the spiking response of significantly responsive neurons to pup call stimuli by doing principal component analysis (PCA) on the peri-stimulus time histograms (PSTHs). Specifically, for each individual neuron, we calculated the mean firing rate across trials with 1 ms bins from the onset of the stimulus to 40 ms after the offset of the stimulus over all trials (120 repetitions for the pup call stimuli). We also estimated the baseline firing rate by calculating the mean firing rate of each neuron within 400 ms to 1000 ms of each stimulus presentation. We then subtracted the baseline firing rate for all significant neurons and concatenated the resulting response over time to get a population response matrix (number of neurons x number of time bins). After normalization (dividing each neuron’s response by its Euclidean norm), we performed PCA on the baseline subtracted and normalized population response matrix. Then we estimated the least number (n) of principal components that can explain 90% variance and performed tSNE with the first n principal components. We used k-means clustering on the high-dimensional tSNE space to cluster the response. The selection of the number of clusters and the quality of the clusters was verified by silhouette values/plots and the plot of total within-cluster sum of squares (WSS).

### STRFs

We estimated the STRFs of neurons using the STAs calculated from the spiking response to the dynamic random chord stimulus described above, as was done in our previous study (Ito et al. 2020; Si et al. 2022).

To construct the data matrix for clustering, we first determined the significant “pixels” in all STRFs. For each neuron, the STRF consists of a 48 (sound frequencies) x 10 (5 ms time bins) x 2 (contra-vs ipsilateral) matrix, which can be displayed as two heatmaps (contra- and ipsilateral), each consisting of 48 x 10 pixels. Each significant neuron has significant pixels (p < 0.001) and non-significant pixels; when examining all significant neurons, we only selected the pixels that were significant in more than 3 neurons. This led to a total of 153 significant pixels.

We then concatenated each neuron’s 153 significant pixels in a single row, to create a 153 by 190 (number of neurons with a significant STRF) matrix as the data matrix for clustering analysis. Then we did PCA on the clustering matrix and used the first 29 PCs that explained 90% variance of the data. We then used k-means clustering methods on the high-dimensional tSNE space of the first 29 PCs, and the quality of the clusters were verified by silhouette values/plots and the plot of total within-cluster sum of squares (WSS).

To identify the resulting clusters of the STRF clustering, we first calculated the average within-ear cosine similarity index (SI) of each STRF cluster; the one with the highest SI value was identified as the “spectral cue neurons” cluster, and the one with the lowest SI value was identified as the “ILD neurons” cluster. To distinguish the rest of the clusters, we then calculated the within-cluster cosine similarity index (cSI) to measure how similar the STRF patterns are within each cluster, and the cluster that has a significantly lower cSI than other clusters contain the highest heterogeneity and was therefore identified as the “others” cluster. Finally, between the remaining two clusters, we calculated the average response latency and the one with a significantly shorter latency was identified as the “fast neurons” cluster, and the other cluster was then identified as the “slow neurons” cluster. We did the same procedure of identifying these STRF clusters in the exploratory dataset and the blinded dataset, and the results turned out to be consistent between the two halves of the full dataset.

## Clustering naturalistic response with STRF clusters

To statistically estimate how well the STRF clusters can be used to distinguish the response of SC neurons to all auditory stimuli tested, we used a bootstrap approach to estimate how well separated the clustered responses grouped by STRF labels are. We first used the 5 STRF labels (spec. ILD, fast, slow, and others) to group the neurons that are significantly responsive to both random chord and another stimulus (original pup call, pup call stretched 1.25 times, pup call stretched 1.25^2^ times, pup call stretched 1.25^3^ times, 60 ms chirp that goes from 20 to 80 kHz, 20 ms chirp that goes from 20 to 80 kHz, 10 ms chirp that goes from 20 to 80 kHz, 10 ms chirp that goes from 80 to 20 kHz). For each stimulus, with these defined groups, we calculated the silhouette values (the cosine similarity difference between each neuron and other neurons in the same group versus outside the group) and calculated the average and SEM of such value for neurons that belong to each group and all neurons (black points with error bars in Fig 5C); then, we randomly assigned the labels to neurons and calculated the average and SEM of silhouette values for each group and all neurons, which we repeated 10000 times to get a distribution of the silhouette value for each group and all neurons combined (colorful distributions in Fig 5C). From the relative position of the STRF grouping results to the randomized distribution, we were able to estimate how the STRF cluster labels can significantly separate the response patterns of SC neurons to various auditory stimuli.

